# Nevermore: Target-Conditioned Protein–Ligand Representation Learning for Multi-Objective Lead Optimization with Database-Grounded Retrieval

**DOI:** 10.64898/2026.01.20.700610

**Authors:** Mohammad Saleh Refahi, Milad Toutounchian, Bahrad A. Sokhansanj, Hyunwoo Yoo, James R. Brown, Hai-Feng Ji, Gail Rosen

## Abstract

Target-conditioned molecular design requires optimizing binding affinity to a proposed therapeutic protein target while balancing competing developability constraints (e.g., absorption, distribution, metabolism, excretion, and toxicity; ADMET). Yet many computational pipelines either optimize a single objective or rely on fully de novo generation that can be difficult to control and interpret. We present Nevermore, a target-conditioned, database-grounded framework that combines a geometry-aware protein–ligand affinity oracle with Pareto-aware multi-objective search over an explicit molecular feature space. A central design choice is to optimize in *count-based Morgan fingerprint* space, where each feature corresponds to a chemically meaningful substructure count, enabling discrete, interpretable “bucket-level” edits. Nevermore learns target-conditioned scores by aligning protein and ligand representations under contrastive objectives and using a similarity-based prediction head; the resulting affinity oracle improves over previously reported benchmark baselines, providing a stronger scoring signal for downstream optimization. Nevermore then steers candidate selection by proposing sparse fingerprint edits, re-ranking candidates under multiple objectives, and projecting edited fingerprints back to valid molecules via nearest-neighbor retrieval from a large compound library. This yields efficient screening without exhaustive enumeration and provides transparent attributions that connect optimization steps to concrete chemical motifs. We evaluate Nevermore on two target case studies (Menin and SARS-CoV-2 Mpro). Across targets, the closed-loop search consistently retrieves candidate sets with improved affinity–property trade-offs compared with random sampling and similarity-only retrieval baselines, while maintaining explicit control and interpretability through discrete feature-space edits. These results support database-grounded, feature-space steering as a practical route to target-conditioned multi-objective lead refinement without relying on fully de novo generation.

## 1 Introduction

Modern drug discovery relies on accurate protein–ligand binding affinity prediction to prioritize potential drug candidate compounds before costly synthesis and experimental assays. Deep learning methods, including structure-aware scoring and complex-prediction models, have improved predictive performance in a range of settings [1, 2]. Nevertheless, potency alone is rarely sufficient for translational success. Drug candidates continue to exhibit substantial attrition in development, and late-stage failures are frequently associated with safety liabilities or inadequate efficacy, with pharmacokinetic limitations and toxicity remaining persistent contributors [3, 4].

These challenges have motivated pipelines that jointly optimize potency-related objectives while explicitly accounting for ADMET (absorption, distribution, metabolism, excretion, and toxicity) constraints [5, 6]. Recent inverse-QSAR (Quantitative Structure-Activity Relationship) and multi-objective optimization frameworks operationalize this goal by searching for representation-level changes that improve predicted efficacy while maintaining drug-likeness and safety profiles [7, 8]. In parallel, descriptor- and fingerprint-based approaches have shown that well-chosen molecular representations can remain competitive in data-constrained regimes, especially when coupled with sample-efficient black-box optimization over large libraries [9, 10]. Taken together, these lines of work motivate hybrid strategies that combine fast, interpretable QSAR-style representations with constraint-aware optimization to support target-driven hit and lead refinement[11].

As illustrated in Figure 1, we present **Nevermore**, a target-conditioned framework that bridges binding-affinity modeling with multi-objective refinement via feature-space steering and database-anchored retrieval. We make four contributions:

**Figure 1.**
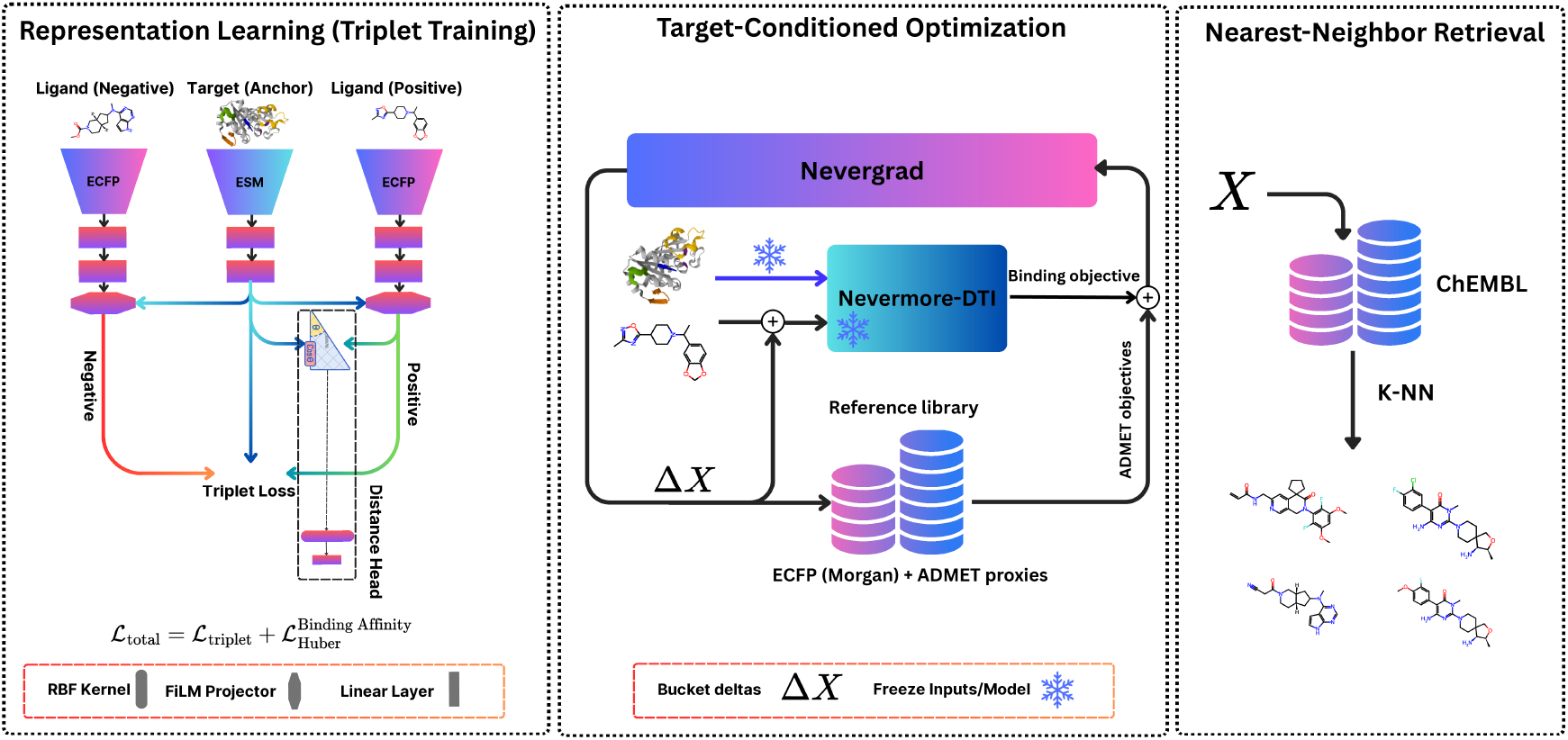
Nevermore pipeline. (Left) Representation learning. We train a geometry-aware DTI oracle with triplet learning to align protein embeddings (ESM) and ligand features (ECFP/Morgan counts) in a shared space using a FiLM projector and a distance-based (RBF) head. **(Middle) Target-conditioned optimization with projection-in-the-loop**. Given a target protein and a baseline ligand fingerprint *X*, Nevergrad proposes sparse integer bucket edits Δ*X*; the DTI oracle is kept frozen (snowflake) while a multiobjective score combines the binding objective with ADMET violation penalties from a reference library). **(Right) Nearest-neighbor retrieval**. Each edited fingerprint is projected back to *valid* molecules via *K*-NN search in a large compound library (e.g., ChEMBL), yielding synthesizable candidate ligands.

### 1. Geometry-aware, metric-aligned protein–ligand representation learning for affinity scoring

We introduce *Nevermore–DTI*, a lightweight target-conditioned oracle that conditions ligand features on the target via a FiLM projector [12], aligns protein–ligand embeddings with contrastive learning under cosine geometry, and predicts affinity with an RBF-style similarity head.

### 2. Target-conditioned, multi-objective derivative-free fingerprint steering

We perform sparse, integer-constrained edits in count-based Morgan fingerprint space using Nevergrad [13], optimizing affinity jointly with developability objectives via explicit constraint penalties (e.g., ADMET proxies).

### 3. Library-anchored, decoding-free realization

We convert optimized fingerprints into concrete candidates by retrieving nearest neighbors from large reference libraries (e.g., ChEMBL), avoiding ambiguous inversion to de novo structures.

### 4. Interpretability and traceability by construction

Because optimization proceeds through sparse feature edits grounded by retrieved neighbors, Nevermore supports direct inspection of which fingerprint dimensions change and how they translate into specific chemical motifs and candidate ligands.

## 2. Related Work

### Protein–ligand affinity prediction and joint embedding models

Protein–ligand affinity prediction depends critically on how proteins and small molecules are represented, and has driven a progression from end-to-end sequence models to pretrained joint-embedding approaches. Early sequence-based predictors encoded protein sequences and ligand SMILES with convolutional networks and regressed affinity from fused features (e.g., DeepDTA) [14], with subsequent variants refining the convolutional interaction modeling [15]. A parallel line of work represents ligands as molecular graphs and couples graph encoders with protein sequence encoders to learn end-to-end interaction predictors (e.g., GNN-CPI) [16]. More recently, transformer architectures improved cross-modal modeling by applying attention over SMILES and protein sequences; MolTrans is a representative example that learns interaction-aware features from both modalities [17].

A complementary viewpoint treats DTI(drug–target interaction) as *representation alignment* : proteins and ligands are first embedded with strong pretrained encoders, then lightweight projection heads are trained so that interacting pairs are close in a shared metric space while mismatched pairs are separated. Protein language models such as ESM [18] and ProtTrans-style models [19] provide transferable protein features, while molecular foundation models such as MolE [20] and SMILES-pretrained transformers such as Chem-BERTa [21] provide general molecular representations. Contrastive and metric-learning objectives explicitly shape this geometry often improving robustness by enforcing relative structure (“which pairs should be closer”) rather than relying only on pointwise regression [22–25]or inexpensive ligand descriptors such as circular (ECFP/Morgan) fingerprints can complement learned embeddings, providing a compact and effective representation for large-library prioritization and retrieval [11, 22, 23].

A complementary viewpoint treats DTI as *representation alignment* : proteins and ligands are embedded with pretrained encoders, and lightweight projection heads are trained so that interacting pairs are close in a shared metric space while mismatched pairs are separated. Protein language models such as ESM [18] and ProtTrans style models [19] provide transferable protein features, while molecular foundation models such as MolE [20] and SMILES pretrained transformers such as ChemBERTa [21] provide general molecular representations. Contrastive and metric learning objectives shape the geometry of this space by enforcing relative similarity constraints, rather than relying only on pointwise regression [22–24]. Within this paradigm, ligand representations vary across works: some use learned molecular embeddings, while others pair protein embeddings with circular fingerprints such as ECFP or Morgan fingerprints [11, 22, 23]. This joint embedding formulation is attractive in practice because encoders can be frozen, representations can be precomputed, and affinity scoring can be implemented with small trainable heads [22, 24].

### Multi objective lead optimization

Lead optimization is inherently multi-objective: candidates must improve affinity while also satisfying development constraints such as solubility, permeability, toxicity risk, and feasibility of synthesis. A common formulation is goal directed optimization, where a proposal mechanism is coupled to fast surrogate evaluators, often including learned ADMET predictors [6].

One influential family of methods performs optimization in a learned continuous latent space. Gómez Bombarelli et al. trained an encoder and decoder to map discrete molecular representations into a continuous vector space, together with property predictors defined on that space [26]. This enables molecular search through sampling and local perturbations around known molecules, as well as gradient based updates in latent space to improve target properties before decoding back to discrete structures [26]. Related generative directions include reinforcement learning over SMILES strings [27, 28] and graph based construction or editing constrained by chemical validity [29]. Evolutionary approaches are also widely used, including fragment based recombination and genetic operators that iteratively refine candidates under multiple objectives [30].

Because optimization can exploit imperfections in surrogate objectives, benchmarking has emphasized validity, diversity, and distribution shift under proxy driven search. GuacaMol provides standardized goal directed tasks and highlights common failure modes when optimizing against approximate property or similarity objectives [31]. In target conditioned pipelines, docking or physics inspired scoring can add structural signal, but it is typically more expensive than learned or descriptor based surrogates and is therefore harder to use as a tight inner loop objective[30].

An alternative direction frames molecular design as black-box optimization over engineered feature spaces. In this setting, descriptors and fingerprints provide explicit, fixed-length representations that can be edited and evaluated efficiently, enabling uncertainty-aware surrogate modeling and sample-efficient Bayesian optimization. Multi-objective variants use Pareto-aware acquisition functions to explore trade-offs directly in objective space, and empirical studies have reported advantages over fixed-weight scalarization baselines in evaluation-limited regimes (i.e., when each evaluation is expensive and only a small number of queries can be afforded), including improved Pareto coverage and diversity [32]. Beyond acquisition design, recent work improves practicality by restricting optimization to adaptively selected subspaces over large descriptor libraries, which can improve data efficiency and interpretability [9]. Related directions also couple proposal mechanisms with multi objective guidance, including LLM guided frameworks for iterative lead improvement [33]. These feature space formulations are particularly compatible with target conditioned pipelines, where affinity is optimized jointly with developability objectives, a setting that we explicitly address in our framework.

## 3 Methods

### 3.1 Affinity oracle

#### Architecture

As the scoring oracle in Nevermore, we use a geometry-aware DTI model following Firm– DTI [23]. Proteins are encoded with a frozen ESM2 protein language model (mean-pooled sequence embedding), and ligands are encoded as RDKit *count* -based Morgan fingerprints (radius *r* = 3, *B* = 1024 bins; sensitivity analysis is reported in the Appendix). Each modality is passed through a learned projection MLP into a shared *d*-dimensional latent space. We denote the projected protein (target) embedding by *z*_*t*_ ∈ ℝ^*d*^ and the projected ligand (drug) embedding by *z*_*d*_ ∈ ℝ^*d*^.

To model target-conditioned interactions, we FiLM-condition the ligand embedding on the protein embedding [12]:

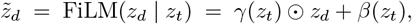

where *γ*(·), *β*(·) : ℝ^*d*^ → ℝ^*d*^ are learned linear maps that produce element-wise scaling and shifting vectors, and ⊙ denotes element-wise multiplication. We *ℓ*_2_-normalize the resulting embeddings and compute a cosine distance,

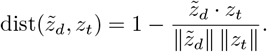

Finally, affinity is predicted via an RBF expansion over this distance with *k* centers 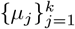 (evenly spaced in [0, 2]),

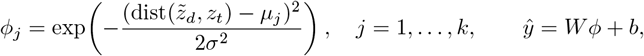

where *ϕ* = [*ϕ*_1_, …, *ϕ*_*k*_] and *ŷ* is the predicted log-affinity.

#### Datasets

The oracle is trained on TDC DTI-DG drug–target pairs [34], using experimentally reported affinity labels (e.g. *pK*_*d*_, *pIC*_50_, or *pXC*_50_). For triplet construction, positives are chosen as higher-affinity compounds for a given protein, and negatives as lower-affinity or cross-target compounds.

#### Training

We train the oracle once, offline with AdamW using the combined objective

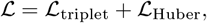

where ℒ_triplet_ enforces a margin between positive and negative pairs in embedding space and _Huber_ regresses *ŷ* onto the experimental log-affinity labels. Full details of the training protocol, hyperparameter settings, and dataset are provided in Appendix. The best checkpoint is selected by validation correlation and then frozen; in all Nevermore experiments the model is used purely as a black-box affinity oracle.

### 3.2 ADMET predictor

To characterize drug-likeness and safety-related properties of each ligand, we compute in silico ADMET profiles using the ADMET-AI [6] platform, a machine learning system that combines graph neural networks with physicochemical descriptors to predict a wide panel of absorption, distribution, metabolism, excretion, and toxicity endpoints for small molecules. For every SMILES string in our training, validation, and retrieval pools, we run ADMET-AI once offline and store the resulting endpoints in a tabular database keyed by a unique molecule identifier.

In this work we focus on a small subset of ADMET-related endpoints that are commonly used in medicinal chemistry: predicted hERG blockade probability (cardiotoxicity risk), predicted human intestinal absorption (HIA Hou), and standard physicochemical/drug-likeness descriptors including logP (lipophilicity), molecular weight, quantitative estimate of drug-likeness (QED), and a Lipinski-rule summary score. To avoid unnecessary reliance on learned predictors, we distinguish *structure-computable* descriptors from *learned* ADMET endpoints: molecular weight, logP, QED, and Lipinski-style summaries are computed deterministically from the molecular graph (e.g., via RDKit), whereas hERG and HIA are biological properties that are not directly computable from SMILES and are obtained from ADMET-AI predictions trained on experimental datasets. Accordingly, predicted ADMET values are used as screening proxies for ranking and retrieval (not as groundtruth safety measurements). The resulting ADMET table provides, for each molecule, a fixed-length vector of standardized ADMET features that we can later query during optimization and retrieval, and that we summarize in the case studies to compare baseline ligands against their nearest neighbors in terms of safety and drug-likeness.

### 3.3 Nevergrad feature-space optimization

We use *Nevergrad*[13] as a derivative-free optimizer to search over an editable *fingerprint space* rather than directly modifying molecular graphs. This choice is convenient in our setting because the search variable is discrete (count buckets), the projection step maps edits back to *real* library molecules, and the resulting pipeline is effectively non-differentiable. We parameterize this space with count-based ECFP (Morgan) fingerprints, which capture local chemical environments in a fixed-length vector amenable to discrete steering and are a widely used, competitive baseline in molecular property modeling compared to coarser structural keys (e.g., MACCS) or more expensive alternatives.[9] While hashed fingerprints can be coarse and non-invertible due to collisions, Nevermore avoids explicit decoding by grounding edits through database retrieval.

Given a target protein sequence *t* and a candidate ligand *s*, our frozen DTI oracle returns a predicted binding affinity *ŷ*(*t, s*). In addition, we associate each library compound with precomputed auxiliary quantities (e.g., its Morgan count fingerprint and developability proxies such as ADMET endpoints). During optimization, once Nevergrad proposes a steered fingerprint, mapping it to candidate molecules and evaluating constraint penalties can be implemented efficiently using nearest-neighbor retrieval followed by lookup of the stored properties for the retrieved compounds.

#### Decision variables and edited fingerprint

Let *X*(*s*) ∈ ℕ^*B*^ denote the count-based Morgan fingerprint of ligand *s* with *B* buckets. Starting from a baseline ligand *s*, we select a small editable set of buckets *S* ⊂ {1, …, *B*} with |*S*| = *m* ≪ *B*. Nevergrad proposes an edit vector ***δ*** ∈ ℝ^*m*^, which we discretize by rounding and enforcing non-negativity:

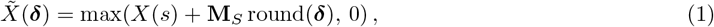

where **M**_*S*_ ∈ {0, 1} ^*B×m*^ is a fixed insertion/selection matrix that places the *m* editable coordinates (indexed by *S*) into the full *B*-dimensional fingerprint, and Δ = round(***δ***) ∈ ℤ^*m*^ are the integer bucket-count adjustments (which may be negative).

#### Projection-in-the-loop (top-*K* neighbor evaluation)

Because 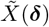 may not correspond to a valid molecule, we project each proposal onto real ligands in a reference library 𝒟 (e.g., ChEMBL) by retrieving the top-*K* nearest neighbors under a weighted *ℓ*_1_ distance computed over the editable buckets:

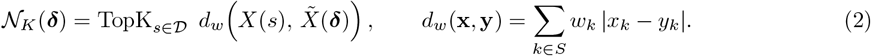

We set *w*_*k*_ = 1 for buckets that are not edited and upweight buckets that Nevergrad changes, so that projection prioritizes agreement on the modified parts of the fingerprint.

#### Joint objective with affinity loss and ADMET penalties

For each retrieved neighbor *s* ∈ 𝒩 _*K*_(***δ***), we evaluate an affinity term from the oracle and an ADMET violation penalty using library endpoints. Let *y*^⋆^ denote the target affinity used in optimization, and let *a*_*j*_(*s*) be the library value of ADMET endpoint *j* ∈ 𝒜 for compound *s* with bounds (*ℓ*_*j*_, *u*_*j*_) and weight *w*_*j*_. We define the per-molecule loss

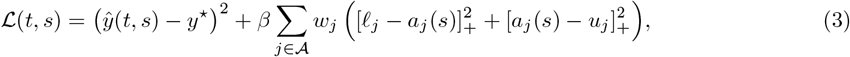

where [*x*]_+_ = max(*x*, 0) and *β* controls the trade-off between affinity and ADMET feasibility. We then use the mean loss over the projected set as the Nevergrad objective,

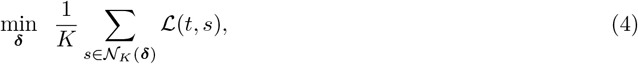

#### Retrieval library

For projection and attribute lookup, we use a ChEMBL-derived[35] retrieval set containing ∼ 2.9 million compounds; all fingerprints and ADMET endpoints are precomputed once offline for fast nearest-neighbor queries.

## 4 Results

### 4.1 DTI oracle model for binding-affinity optimization and interaction prediction

#### Binding-affinity oracle (regression)

We train a geometry-aware protein–ligand affinity predictor to serve as the scoring oracle in Nevermore. We evaluate on the BindingDB Patent benchmark (TDC) [34] using Pearson correlation coefficient (PCC) between predicted and experimental affinities. The oracle encodes the target protein with a frozen protein language model (PLM) and represents ligands using a count-based Morgan fingerprint (ECFP-style), which is then projected into a shared latent space.

Figure 2 ranks methods by PCC. A MolE-based ligand encoder with the same interaction architecture achieves the strongest correlation, while our Morgan-fingerprint variant remains competitive with other PLM-based baselines (approximately 2.5% lower PCC than the MolE variant). We nevertheless use the Morgan-fingerprint oracle in Nevermore because it exposes a *directly editable* discrete feature space (fingerprint buckets), enabling efficient Nevergrad search and straightforward projection back to valid library molecules.

**Figure 2.**
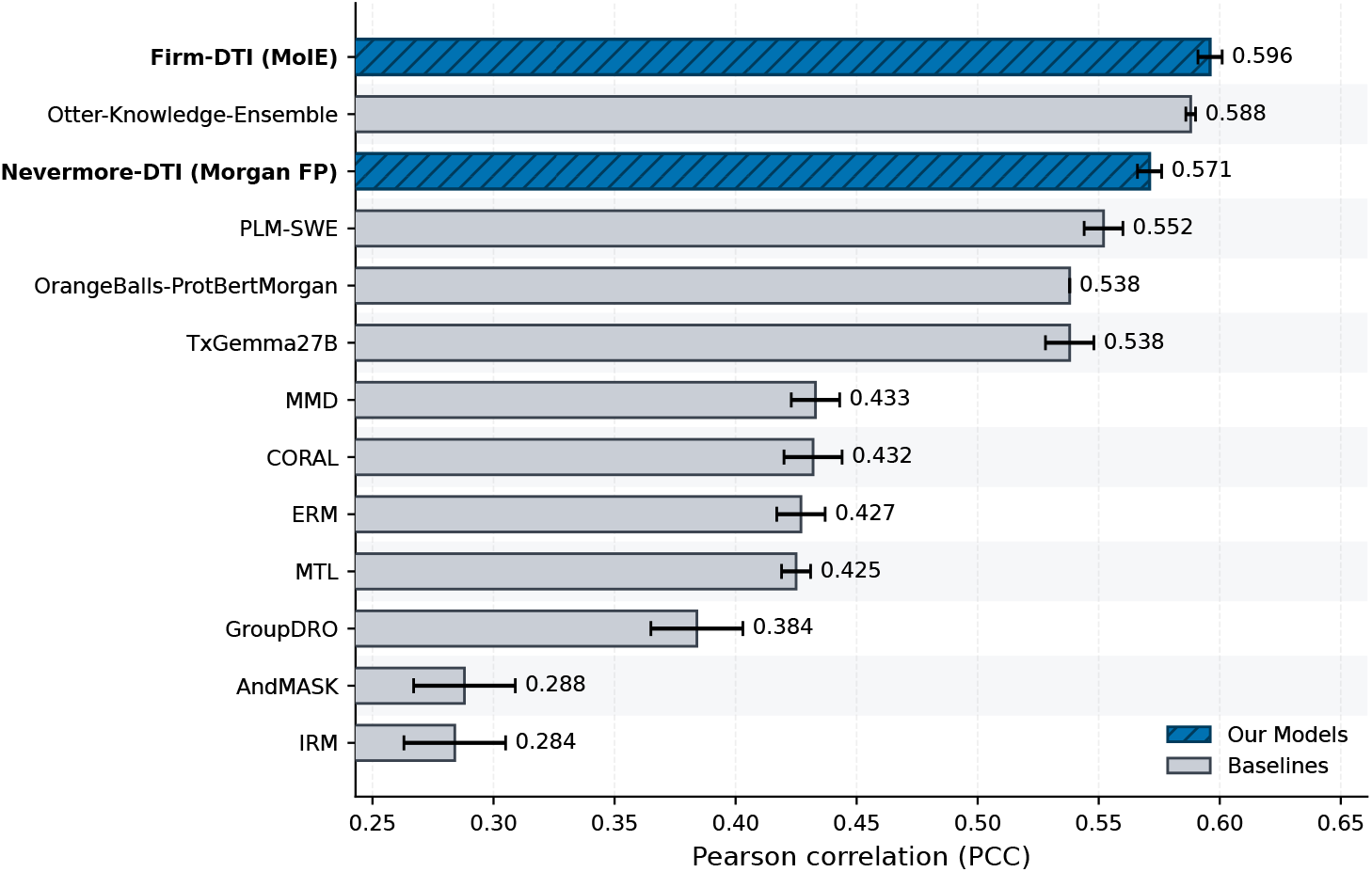
BindingDB Patent (TDC) affinity prediction. Ranked comparison by Pearson correlation (PCC) between predicted and experimental binding affinity. Firm–DTI (MolE) achieves the highest PCC, followed by Otter-Knowledge-Ensemble and Nevermore–DTI (Morgan FP).

#### DTI classification (binary interaction)

In addition to affinity regression, we train the same geometry-aware protein–ligand model as a binary drug–target interaction (DTI) classifier by replacing the regression head with a sigmoid classification head and optimizing cross-entropy loss [23]. We evaluate on three standard benchmarks DAVIS [36], BindingDB [37], and BIOSNAP (ChG-Miner) [38] reporting AUROC and AUPR (mean ± s.e.m. across seeds). Following MolTrans [17], we binarize DAVIS and BindingDB using *K*_*d*_ < 30 as positive and *K*_*d*_ ≥ 30 as negative.

Table 1 summarizes results for two ligand representations within this architecture: a MolE-based ligand encoder and an editable count-based Morgan fingerprint (Morgan FP). Across datasets, the MolE variant yields the strongest overall performance, while the Morgan-FP variant remains competitive and typically ranks second, outperforming or matching several established baselines.

**Table 1:** Comparison on BIOSNAP, BindingDB, and DAVIS datasets. Mean ± s.e.m. over multiple seeds.

| Benchmark | Model | AUPR | AUROC |
| --- | --- | --- | --- |
| BIOSNAP (19,238) | **Firm-DTI (MolE)** | **0.919 ± 0.001** | **0.910 ± 0.001** |
| | Nevermore-DTI (Morgan FP) | 0.890 ± 0.006 | 0.891 ± 0.006 |
| | MolTrans[17] | 0.885 ± 0.005 | 0.876 ± 0.007 |
| | GNN-CPI[16] | 0.890 ± 0.004 | 0.879 ± 0.007 |
| | DeepConv-DTI[15] | 0.889 ± 0.005 | 0.883 ± 0.002 |
| BindingDB (12,668) | **Firm-DTI (MolE)** | **0.647 ± 0.001** | **0.916 ± 0.001** |
| | Nevermore-DTI (Morgan FP) | 0.591 ± 0.007 | 0.903 ± 0.008 |
| | MolTrans[17] | 0.598 ± 0.013 | 0.898 ± 0.009 |
| | GNN-CPI[16] | 0.578 ± 0.015 | 0.900 ± 0.004 |
| | DeepConv-DTI[15] | 0.611 ± 0.015 | 0.908 ± 0.004 |
| DAVIS (2,086) | **Firm-DTI (MolE)** | **0.454 ± 0.001** | 0.870 ± 0.001 |
| | Nevermore-DTI (Morgan FP) | 0.437 ± 0.003 | **0.920 ± 0.002** |
| | MolTrans[17] | 0.335 ± 0.017 | 0.889 ± 0.007 |
| | GNN-CPI[16] | 0.269 ± 0.020 | 0.840 ± 0.012 |
| | DeepConv-DTI[15] | 0.299 ± 0.039 | 0.884 ± 0.008 |

### 4.2 Sensitivity to objective weight, retrieval metric, and edit budget

Nevermore optimizes sparse edits to Morgan *count* buckets with Nevergrad, while staying *database-grounded* via projection-in-the-loop: after each edit, we retrieve the top-*K* nearest library molecules and summarize the retrieved set by predicted affinity and precomputed ADMET endpoints. We report (i) ΔAffinity (vs. the baseline ligand), (ii) Mol Aff (mean affinity over the retrieved top-*K*), and (iii) mean ADMET endpoints over the same set (MW, logP, hERG risk, QED; sign-flipped where needed so higher is better). For comparison across metrics, we z-normalize columns across runs and visualize sweep heatmaps, with stars marking the best configuration(s) per metric. We evaluate 10 protein targets held out from our training, and for stable statistics we summarize *K*=100 retrieved molecules per target (1000 total).

#### Hyperparameter sweeps and trade-offs

Figure 3 summarizes four controlled sweeps: (i) multi-objective weight *β*, (ii) retrieval distance (Jaccard vs. weighted *ℓ*_1_), (iii) optimization budget (Nevergrad iterations), and (iv) number of editable buckets.

**Figure 3.**
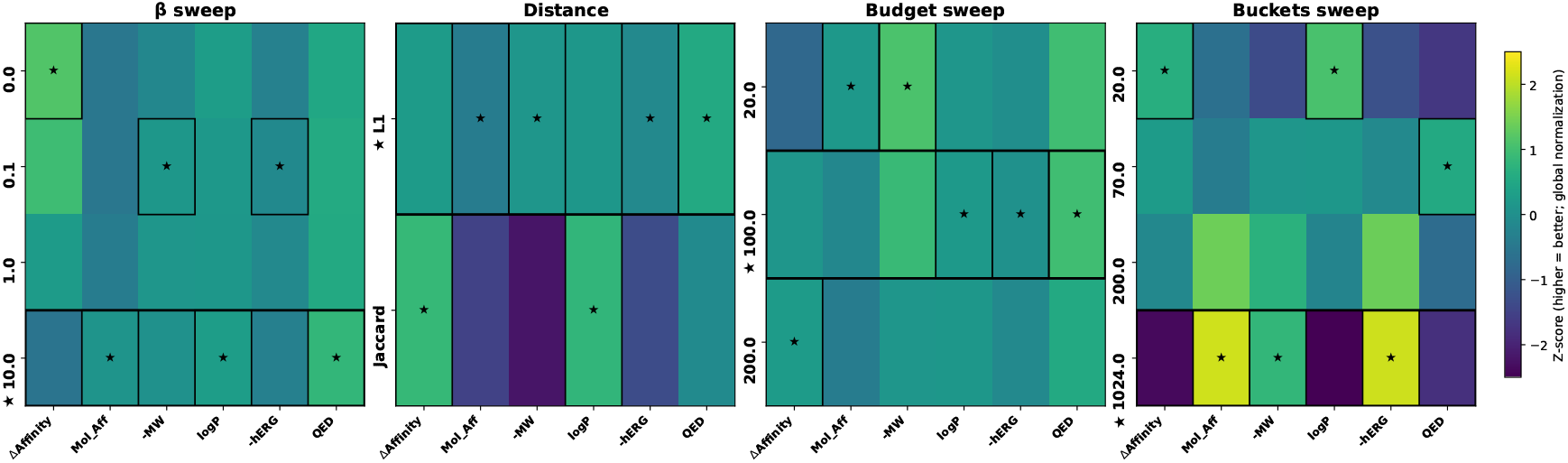
Optimization dynamics across objectives and search settings. Each panel varies one factor (left-to-right: *β*, distance, budget, editable buckets). Columns show z-normalized metrics (higher is better): ΔAffinity, Mol Aff, −MW, logP, −hERG, QED. Stars mark best configuration(s) per column within each panel.

Increasing *β* shifts the search from affinity-only toward constraint-aware optimization, revealing a consistent trade-off: low-*β* settings tend to maximize ΔAffinity, whereas higher *β* improves the *projected* neigh-borhood (higher Mol Aff and QED) and reduces liability-aligned endpoints such as MW and hERG risk (after sign-flipping). Across distances, weighted *ℓ*_1_ retrieval produces more stable and higher-quality neighborhoods than Jaccard, consistent with magnitude-aware matching being better aligned with Morgan-count edits. Increasing the iteration budget generally improves ΔAffinity, but retrieved-set ADMET/QED often peak at moderate budgets, suggesting diminishing returns once the optimizer begins exploiting feature edits that project less reliably. Finally, enlarging the editable bucket set |*S*| increases expressivity but also raises the risk of off-library edits: small |*S*| can yield the best ΔAffinity, while intermediate-to-large |*S*| often improves neighborhood statistics, with QED frequently peaking at intermediate sizes—too few buckets under-express multi-property adjustments, while too many can induce unstable or compensatory edits that projection cannot consistently realize.

Overall, the sweeps highlight that feature-space gains can decouple from database-grounded outcomes: Nevergrad may propose bucket combinations that are rare or absent in the library (off-manifold), and nearest-neighbor projection can be discontinuous, so small edits can abruptly change the retrieved set. As a result, configurations that maximize ΔAffinity do not necessarily yield the best Mol Aff or ADMET trends in the projected neighborhood.

### 4.3 Case Study: Cross-target evaluation of multi-objective ligand optimization

#### 4.3.1 Target selection and baseline ligands

We study ligand optimization under competing objectives on two structurally distinct targets to stresstest generality across pocket geometry and chemical priors: **Menin** (oncology-relevant, PPI-like pocket; Menin inhibitors primarily disrupt Menin–protein interactions rather than inhibiting an intrinsic enzymatic activity) [39, 40] and **SARS-CoV-2 M**^**pro**^ (viral cysteine protease; inhibitors target the catalytic substrate-binding cleft) [41].

##### Menin

As a stringent, clinically grounded starting point, we use **revumenib** (Revuforj)[42] as the baseline Menin ligand. Revumenib is an FDA-approved *menin inhibitor* used in molecularly defined acute leukemias. Starting from an approved, highly optimized scaffold makes Menin a challenging test case: improvements over local similarity neighborhoods are expected to be non-trivial, so any gains in multi-objective trade-offs are especially meaningful.

##### SARS-CoV-2 M^**pro**^

For M^pro^, we use **cinanserin**[43] as the baseline to intentionally begin from a chemically plausible but safety-limited scaffold. PubChem reports an acute oral toxicity warning (H302: harmful if swallowed), which motivates optimization beyond affinity alone and provides a clear stress test for multi-objective decision-making.

For each target, we keep the baseline ligand fixed and generate a fixed budget of candidates (*N* = **100**) from the same ChEMBL-scale library (∼2.9M compounds) using three strategies: (i) **Random** sampling from the library; (ii) **Similarity retrieval** around the baseline using `RDKFP+Tversky`, i.e., we compute RDKit topological fingerprints for all library molecules and retrieve the nearest neighbors to the baseline under the Tversky similarity (an asymmetric generalization of Jaccard/Tanimoto that can emphasize “containment” of baseline substructures); and (iii) **Nevermore**, which performs multi-objective optimization in fingerprint count-space and proposes discrete bucket edits, then projects each proposal back to valid molecules via nearest-neighbor retrieval from the same library. Unless otherwise stated, Nevermore uses 70 bucket edits per run and an *ℓ*_1_ retrieval distance with *β* = 1 to balance affinity against property-based penalties.

Across both targets, Nevermore achieves the most favorable multi-objective trade-off and a more stable physicochemical profile. On M^pro^, it improves over similarity retrieval with lower hERG (0.411 vs. 0.500) and higher QED (0.740 vs. 0.676) while maintaining comparable affinity (7.283 vs. 7.229); it also avoids the extreme variability seen under random sampling (e.g., very large MW variance). On Menin, the gains are larger: Nevermore attains the highest affinity (7.060 vs. 6.934) while substantially reducing hERG (0.454 vs. 0.625) and increasing QED (0.712 vs. 0.545) relative to similarity retrieval.

#### 4.3.2 Interpreting optimized bucket edits: from buckets to fragments and target-specific levers

To make Nevermore mechanistically interpretable, we analyze the *bucket edits* produced by the optimizer. A “bucket” is a group of count-based fingerprint features, and an *edited bucket* is one whose count is increased or decreased relative to the baseline. Because edits occur in fingerprint space and final candidates are realized by nearest-neighbor retrieval, a bucket edit should be read as a *directional preference*: it biases retrieval toward library molecules that contain (or avoid) the corresponding local substructures.

We compare the edited-bucket sets obtained when optimizing each objective in isolation (affinity-only or single-proxy-only) against the set obtained under the full multi-objective score (Fig. 4). Intersections indicate *shared levers*, structural motifs that simultaneously improve multiple objectives. In contrast, non-overlapping regions highlight *objective-specific levers* introduced only when that constraint is active. This set-level view cleanly separates (i) edits that preserve binding signal (typically shared with affinity) from (ii) additional edits required to reduce proxy violations, illustrating why high affinity *alone* is generally insufficient for producing candidates with improved safety/quality profiles.

**Figure 4.**
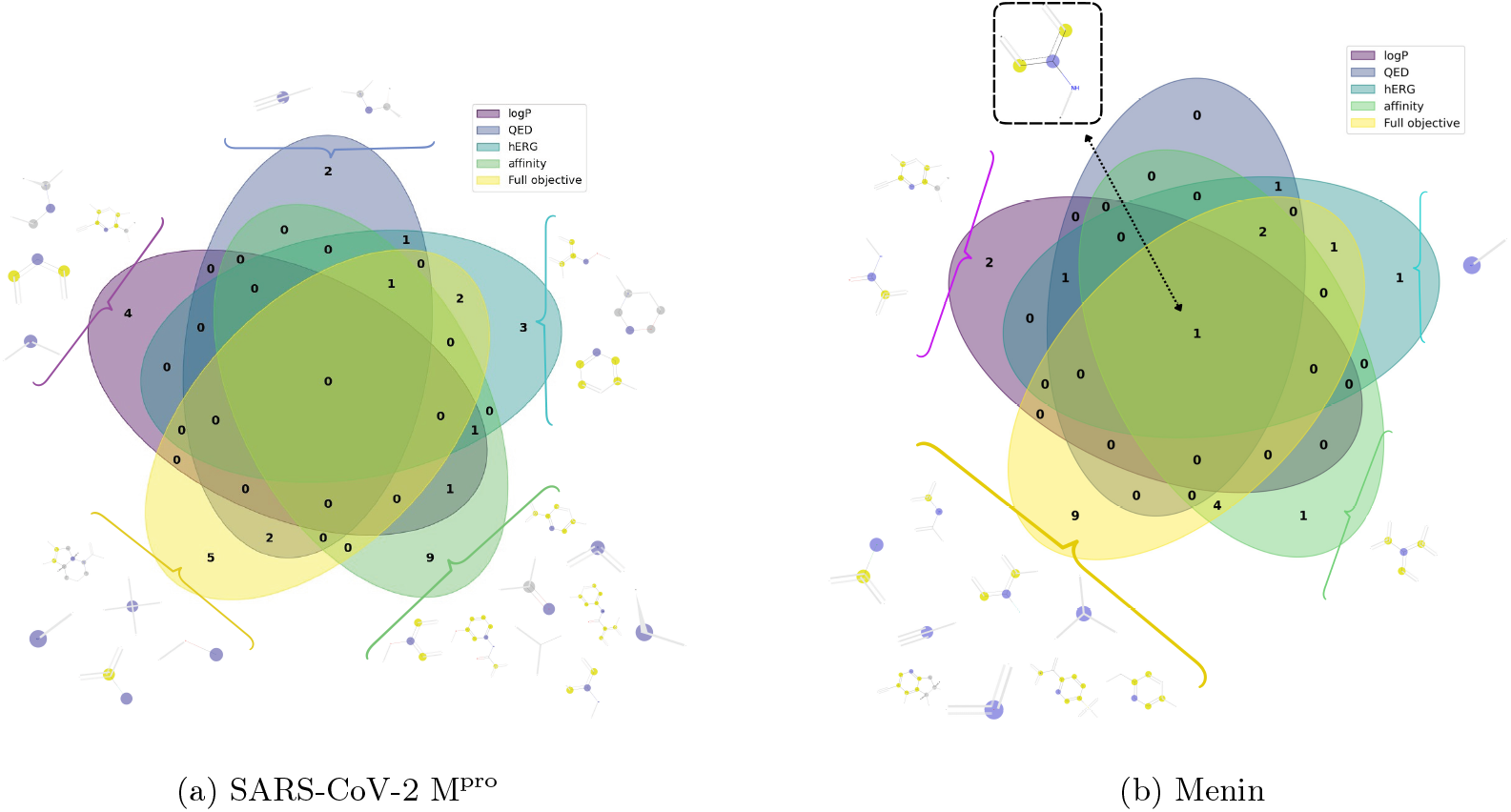
Overlap of optimized bucket edits across objectives. For each target, we show the sets of Morgan-count buckets whose values were adjusted relative to the baseline ligand (Cinanserin for M^pro^, Revumenib for Menin) during objective-specific optimization (affinity or a single ADMET proxy) and during full multi-objective optimization. Intersections highlight *shared* edit directions that improve multiple objectives simultaneously, whereas non-overlapping regions identify *objective-specific* edits that are activated only when a particular proxy constraint is enforced.

To translate bucket indices into chemistry, we map each edited bucket to representative Morgan bits and extract the corresponding atom environments (“bits-to-fragments”). This yields concrete fragment-level hypotheses about *which chemical motifs* Nevermore repeatedly selects under each objective and how these motifs differ by target.

##### Target-specific optimization patterns

For **Menin** (baseline Revumenib), Fig. 4 shows substantial overlap between the objective-specific edit sets and the full multi-objective run, indicating that several edit directions are broadly beneficial rather than proxy-specific. Notably, the *amide*-associated environment emerges as a shared lever across objectives (common to all runs), consistent with carbonyl-containing motifs acting as a stable “core” adjustment that improves multiple scores without strongly violating constraints. Beyond this shared amide-driven core, the full objective introduces additional edits that map to compact polarity and tuning fragments, including *nitrile*-like and *sulfonyl* -like environments, together with *halide*-associated environments. Mechanistically, this pattern supports a “balance” strategy: amide/nitrile/sulfonyl fragments serve as property-control levers (increasing polarity and reducing proxy penalties), while halide-associated edits help preserve or recover affinity by favoring hydrophobic pocket complementarity.

For **M**^**pro**^ (baseline Cinanserin), the decoded bucket edits reveal chemically intuitive, objective-specific levers. hERG-driven optimization is dominated by *neutral polar linker* motifs (ether-like environments), while logP and QED runs preferentially introduce *compact polar* substructures (nitrile/amide/ether) that shift overall polarity without large structural changes. In contrast, affinity-driven runs emphasize *carbonyl H-bond acceptor* environments and scaffold/topology edits that modulate pocket complementarity. The full multi-objective run reconciles these pressures by retaining polarity-increasing edits needed to reduce proxy violations while adding hydrophobic tuning (halide-associated environments) to preserve or recover binding signal, producing a balanced edit set relative to the Cinanserin baseline.

#### 4.3.3 Pareto trade-offs: affinity vs. safety/quality proxies

Multi-objective lead optimization is inherently a *Pareto* problem [32]: improving one objective (e.g., predicted affinity) can degrade others (e.g., safety and physicochemical constraints), so no single metric uniquely defines the “best” candidate. We therefore visualize candidate sets using Pareto-style trade-off plots (Figures 5 and 6) to support decision-making under competing objectives.

**Figure 5.**
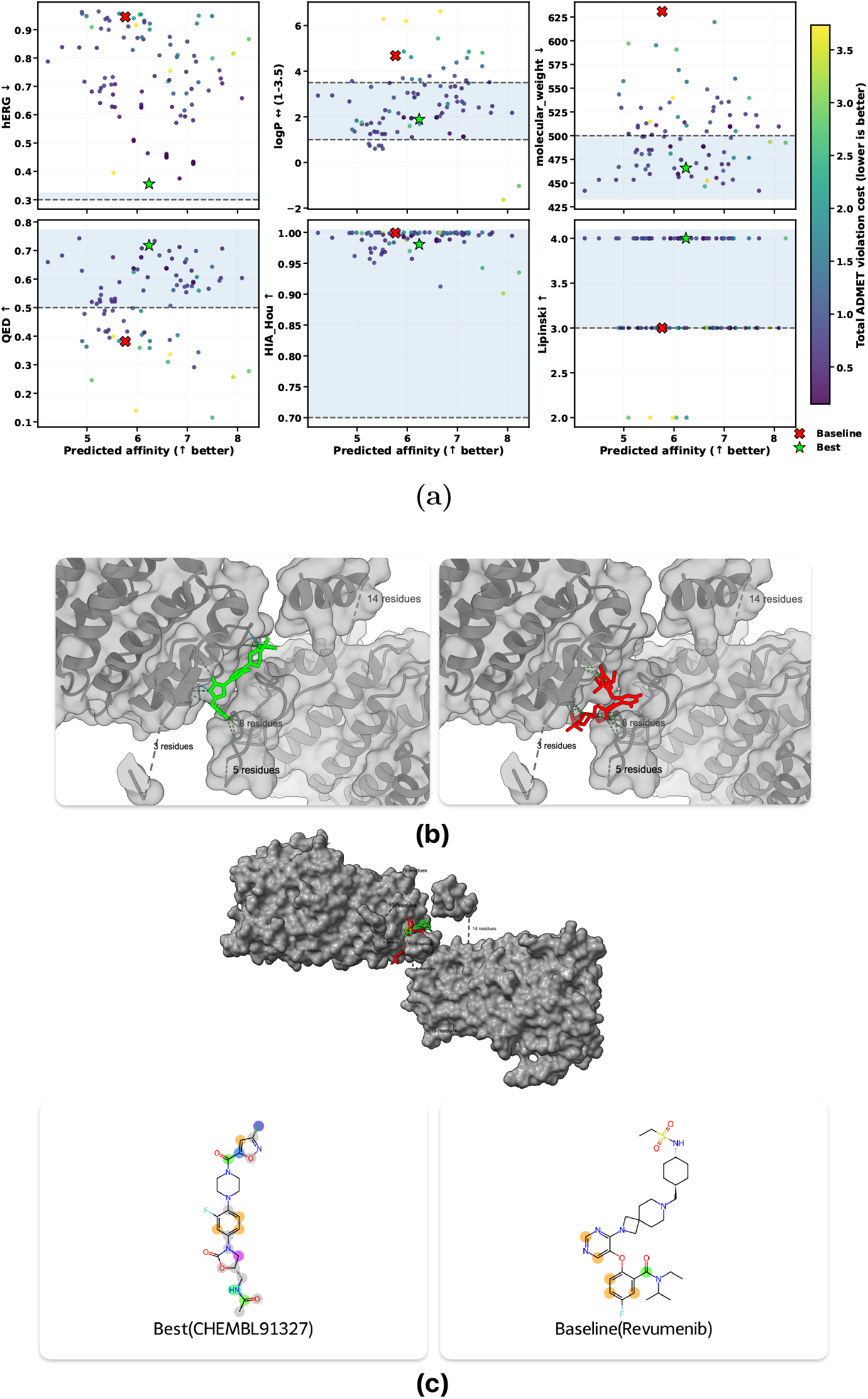
Menin. **(a)** Pareto trade-offs between predicted affinity and ADMET objectives (color: total ADMET-violation cost); baseline Revumenib (red ×) vs selected optimized CHEMBL91327 (green ⋆). **(b)** Pose overlay (baseline red, optimized green) with polar contacts. **(c)** Projection of *adjusted* Morgan-count buckets onto structure (colored centers, gray environm_1_e_4_nts): baseline realizes **2** adjusted buckets vs optimized **7**.

**Figure 6.**
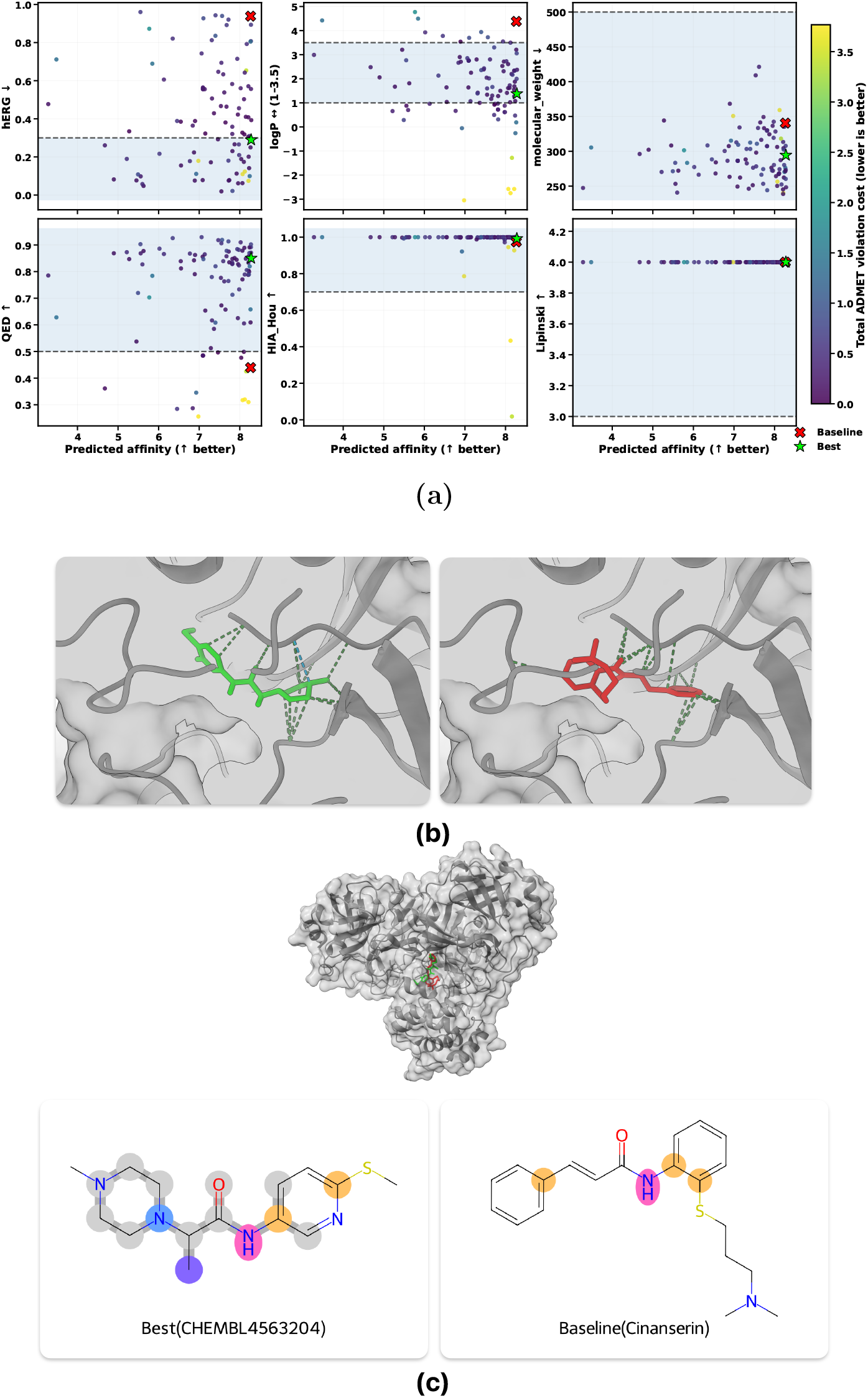
M_pro_. **(a)** Pareto trade-offs between predicted affinity and ADMET objectives (color: total ADMET-violation cost); baseline Cinanserin (red ×) vs selected optimized CHEMBL4563204 (green ⋆). **(b)** Pose overlay (baseline red, optimized green) with polar contacts. **(c)** Adjusted-bucket projection (colored centers, gray environments): baseline realizes **2** adjusted buckets vs optimized **5**.

In each panel, the x-axis is predicted affinity (higher is better) and the y-axis is a single proxy objective (hERG, QED, MW, logP, HIA Hou, or Lipinski). Dashed lines (and shaded regions) indicate the preferred constraint direction or acceptable range (e.g., lower hERG and MW, higher QED, and logP within a target interval). Each point corresponds to one of the *N* = 100 retrieved candidates; points are colored by the *total constraint-violation cost*, summarizing how well a candidate satisfies the full set of proxy constraints. The baseline ligand is highlighted (red ×), and the selected “best” candidate (green ⋆) is the lowest-violation point among high-affinity candidates.

##### Menin

Figure 5 illustrates an even stronger multi-objective shift: the baseline violates multiple constraints (including hERG, MW, and logP), whereas optimized candidates populate regions with higher affinity and lower overall violation. The selected best candidate moves into the feasible ranges for most proxies while improving affinity, indicating that Nevermore can identify meaningful Pareto improvements even for a challenging PPI-like pocket.

##### M^**pro**^

Figure 6 shows that the baseline lies in a high-affinity region but violates key proxy constraints (notably elevated hERG, sub-threshold QED, and out-of-range logP). In contrast, Nevermore shifts candidates toward the feasible (shaded) regions across multiple panels simultaneously: several candidates retain near-baseline affinity while exhibiting markedly lower constraint-violation cost (darker colors), and the selected best point achieves a substantially improved safety/quality profile with minimal affinity compromise. This is consistent with the aggregate improvements reported in Table 2.

**Table 2:** Menin and SARS-CoV-2 M^pro^ candidate-set summary (*N* = 100). Mean ± std over generated candidates (excluding baseline). Best per target is bold.

| Method | Affinity↑ | hERG↓ | QED↑ | MW↓ |
| --- | --- | --- | --- | --- |
| **SARS-CoV-2 M<sup>Pro</sup>** |  |  |  |  |
| Baseline | 8.266 | 0.939 | 0.440 | 340.492 |
| Random | 7.098 ± 1.299 | 0.614 ± 0.285 | 0.526 ± 0.230 | 462.960 ± 453.090 |
| RDKFP+Tversky | 7.229 ± 0.839 | 0.500 ± 0.231 | 0.676 ± 0.203 | 301.715 ± 47.191 |
| Nevermore | **7.283 ± 1.064** | **0.411 ± 0.274** | **0.740 ± 0.179** | **292.959 ± 34.844** |
| **Menin** |  |  |  |  |
| Baseline | 5.765 | 0.945 | 0.382 | 630.831 |
| Random | 6.056 ± 1.362 | 0.858 ± 0.113 | 0.623 ± 0.179 | 449.249 ± 104.242 |
| RDKFP+Tversky | 6.934 ± 0.998 | 0.625 ± 0.284 | 0.545 ± 0.220 | 408.422 ± 128.816 |
| Nevermore | **7.060 ± 0.707** | **0.454 ± 0.230** | **0.712 ± 0.154** | **319.844 ± 72.276** |

**Table 3:** Dataset split sizes used for training and evaluation.

| Dataset | Train | Test | Val |
| --- | --- | --- | --- |
| DAVIS [17] | 2,086 | 6,017 | 3,006 |
| BindingDB [17] | 12,668 | 13,289 | 6,484 |
| BIOSNAP [17] | 19,238 | 5,497 | 2,748 |
| TDC DTI-DG [34] | 146,479 | 49,028 | 36,539 |

#### 4.3.4 Bucket-level attribution of discrete Nevermore edits

After selecting a representative optimized ligand for each target from the Pareto set, we analyze *which fingerprint dimensions were explicitly edited* by Nevermore and where those edits manifest on the chemical graph. Nevermore operates in Morgan-count space with a fixed bucketization: each Morgan feature identifier fid is mapped to a bucket index *b* = fid mod *B*, and the optimizer proposes a sparse adjustment vector Δ**c** over buckets (positive or negative count edits). We focus on the set of *adjusted buckets* 𝒜 = {*b* : Δ*c*_*b*_ ≠ 0} and project them back onto structure using RDKit’s Morgan *bitInfo*. For each adjusted bucket, we extract the corresponding Morgan feature *center atom* (the anchor of the local environment) and its radius-defined neighborhood. In the visualization, colored circles mark center atoms of adjusted buckets (color encodes bucket identity), while gray highlights show the surrounding atoms/bonds that define the corresponding Morgan environments; all other atoms are left unhighlighted. Quantitatively, we report how many adjusted buckets are realized in a molecule (presence of at least one matching center/environment) and how many distinct centers/environments they induce.

##### Menin (Baseline: Revumenib; Optimized: CHEMBL91327)

The baseline ligand realizes **2** adjusted buckets, whereas the optimized ligand realizes **7** adjusted buckets. This is reflected in the number of activated feature centers (**6** in the baseline versus **11** in the optimized ligand) and the extent of structural context supporting those activations (6 environment atoms and 0 bonds in the baseline versus 29 atoms and 18 bonds in the optimized ligand). The optimized candidate therefore exhibits markedly higher coverage of the adjusted bucket set 𝒜 and a richer set of bucket-aligned local environments, providing an interpretable, structure-level explanation for why the Pareto-selected ligand better reflects the optimizer’s discrete fingerprint edits than the baseline.

##### M_pro_ (Baseline: Cinanserin; Optimized: CHEMBL4563204)

The baseline ligand realizes only **2** adjusted buckets (visible as **4** colored center atoms), whereas the optimized ligand realizes **5** adjusted buckets (with **6** colored centers) and induces substantially broader local contexts (20 environment atoms and 15 bonds highlighted). This indicates that the optimized molecule matches a larger fraction of the *intended* discrete edits proposed in bucket space and does so through more extended substructure neighborhoods, consistent with Nevermore introducing/strengthening multiple bucket-aligned motifs rather than relying on a small number of isolated features. Adjusted buckets that appear in both ligands correspond to shared local motifs (similar center placements), while buckets appearing only in the optimized ligand reflect newly introduced or substantially modified motifs relative to the baseline scaffold.

#### 4.3.5 Structural intuition: baseline vs. optimized ligand binding poses

To provide qualitative structural support for the discrete edit directions proposed by Nevermore, we visualized docking poses for each target using the baseline ligand (red) and the Pareto-selected Nevermore-optimized ligand (green). For each receptor, we retain *clash-free* poses under ChimeraX [44] clash detection and compare (i) whether the optimized ligand preserves a plausible anchoring configuration in the binding site and (ii) whether it extends into additional pocket regions or interaction opportunities without introducing steric conflicts. We also overlay baseline and optimized poses to highlight changes in site occupancy. Dashed interaction lines indicate polar contacts (e.g., hydrogen bonds) and are used only for qualitative comparison.

##### Menin (Baseline: Revumenib; Optimized: CHEMBL91327)

For Menin, the optimized ligand remains in the same functional binding region as the baseline but shows a clearer increase in interaction density and site engagement. Under the same contact definition, the optimized pose yields substantially more protein–ligand contacts (baseline: 320; optimized: 515), indicating that the optimized scaffold occupies the pocket more extensively and engages a larger set of surrounding residues. In addition, the optimized ligand forms a markedly stronger polar contact network (9 hydrogen bonds detected for the optimized pose), consistent with enhanced electrostatic complementarity in the binding site. Overall, the Menin poses provide a structural rationale aligned with the Nevermore edit strategy: the optimized candidate maintains a clash-free anchoring configuration while expanding the set of stabilizing local interactions relative to the baseline.

##### M_pro_ (Baseline: Cinanserin; Optimized: CHEMBL4563204)

The overlay indicates that both ligands occupy the same binding cleft with broadly similar pocket packing. Consistent with this visual similarity, the total contact counts are comparable under identical ChimeraX settings (baseline: 396 contacts; optimized: 385 contacts), suggesting similar overall burial and shape complementarity. Importantly, the optimized ligand preserves the core anchoring geometry while exhibiting a slightly richer polar interaction pattern, forming **2** hydrogen bonds versus **1** for the baseline. Together, these observations support the interpretation that Nevermore improves interaction quality (additional stabilizing polar contact) while maintaining a feasible binding pose rather than achieving gains via unrealistic rearrangements.

## 5 Discussion

Nevermore is a fast target conditioned multi objective framework for lead optimization that couples (i) an affinity oracle built from pretrained protein language model embeddings and count based Morgan fingerprints, (ii) derivative free Pareto optimization over sparse discrete fingerprint edits to balance affinity with ADMET constraints, and (iii) projection of edited fingerprints back to real compounds via large library nearest neighbor retrieval. By optimizing a target specific score under multiple objectives while staying anchored to an existing chemical library, Nevermore aims to support practical lead refinement with explicit control over potency and developability. Conceptually, this design sits between deep QSAR and proteochemometrics pipelines, where engineered ligand descriptors remain useful in data limited regimes, and modern target conditioned predictors that leverage pretrained protein representations [11, 14]. Using count based Morgan fingerprints keeps the decision variables discrete and directly editable, which is difficult to achieve with continuous latent molecular embeddings [10].

Compared with goal directed de novo generation, such as latent space models, reinforcement learning, or graph editing, Nevermore prioritizes validity and accessibility by grounding optimization in a curated compound library. This choice helps mitigate common failure modes where proxy optimization drifts away from realistic chemistry, which has been documented in benchmark settings [26–28, 31]. At the same time, feature space optimization aligns naturally with black box search over molecular descriptors and Pareto aware selection, which is particularly relevant when objectives conflict [9, 32].

Empirically, the Morgan fingerprint oracle remains competitive as an affinity predictor and, importantly, exposes the editable feature space required for optimization. In the Menin and SARS CoV 2 M^pro^ case studies, Nevermore shifts candidate sets toward improved trade offs relative to random sampling and baseline centered similarity retrieval, and it supports interpretation through bucket to fragment mappings and qualitative pose inspection. While these analyses are not a substitute for experimental validation, they provide a concrete link between discrete fingerprint edits, recurring substructures, and plausible binding site interactions.

We highlight two limitations that are important for correct interpretation. First, hashed Morgan fingerprints can collide: distinct substructures may map to the same bucket, so an edited bucket can correspond to multiple underlying chemical causes, limiting identifiability of the true structural driver. Accordingly, bucket-to-fragment attributions should be interpreted as *hypothesis-generating* rather than uniquely identifying a single substructure. In our pipeline, this ambiguity is partially mitigated by projecting edited fingerprints back to *real* library molecules via nearest-neighbor retrieval, which anchors edits to concrete, chemically valid candidates. More broadly, recent work demonstrates that ECFP-style fingerprints can be reverse engineered into molecular structures using deterministic enumeration and generative models, suggesting a complementary projection module that could decode optimized fingerprints into candidate structures beyond pure library retrieval [45].

Second, the approach depends on the coverage of the retrieval library and on the calibration of the affinity and ADMET predictors; if either is weak for a given target or chemistry region, the attainable improvements will be limited. These considerations motivate follow up work on collision aware representations and uncertainty aware scoring, as well as tighter integration with experimental feedback.

## 6 Conclusion

Nevermore provides a database grounded approach to target conditioned lead optimization using discrete interpretable fingerprint edits with derivative free multi objective constraints and library retrieval. The main contribution is methodological: it turns QSAR style descriptors into an editable optimization space while maintaining chemical validity by projecting candidates back to real compounds. Across distinct targets, the results support the feasibility of multi objective selection in silico and offer a practical workflow that is fast enough for large library settings, interpretable at the level of fingerprint driven substructures, and compatible with downstream decision making in medicinal chemistry. At the same time, the method can fail when fingerprint collisions or limited library coverage obscure the intended edit directions, highlighting clear opportunities for collision aware and uncertainty aware extensions. Future work that incorporates experimental feedback and improved representations can strengthen reliability and further position this framework as a useful component in iterative lead optimization pipelines.

## Data and Code Availability

All code for Nevermore (including training, optimization, and evaluation scripts) is publicly available at https://github.com/EESI/nevermore.

Benchmark datasets used in this study are publicly available. We obtain the BindingDB Patent and DTI-DG tasks through the Therapeutics Data Commons (TDC) at https://tdcommons.ai/. For the standard in-domain DTI benchmarks (e.g., DAVIS and BindingDB) and their preprocessing/binarization used in MolTrans-style evaluations, we follow the publicly released dataset resources at https://github.com/kexinhuang12345/MolTrans/tree/master/dataset. Any additional preprocessing scripts and experiment configurations used in this work are provided in our repository.

## Acknowledgements

This work is supported by National Science Foundation under Grant Numbers #1936791, #1919691 and #2107108. We thank the University Research Computing Facility for their paid services.

## A. Supplementary Methods

### A.1 Dataset statistics

We use four widely used drug–target interaction (DTI) benchmarks: DAVIS, BindingDB, BIOSNAP (ChG-Miner), and TDC DTI-DG [34, 36–38]. DAVIS, BindingDB, and BIOSNAP follow standard in-domain splits commonly used in prior work. We additionally include TDC DTI-DG, which is designed for out-of-domain (cold-start) evaluation with novel targets and/or compounds. In our pipeline, models trained on TDC DTI-DG are used as the affinity-scoring oracle for constrained optimization, and we also report their predictive performance on the corresponding affinity task.

### A.2 Training details

We trained our model (Nevermore–DTI) on a single NVIDIA H100 GPU with 80 GiB of memory. The total number of parameters is ≈3.5 × 10^9^, with the vast majority residing in the pretrained encoders. During training, we update only ∼ 9 million parameters, while keeping the pretrained encoders frozen. For the protein encoder, we use the ESM2 model `esm2 t12 35M UR50D` from Facebook AI’s repository^1^.

We optimize with AdamW (*ϵ* = 10^−6^) and use a cosine learning-rate schedule with linear warmup. The main hyperparameters are summarized in Table 4.

**Table 4:**
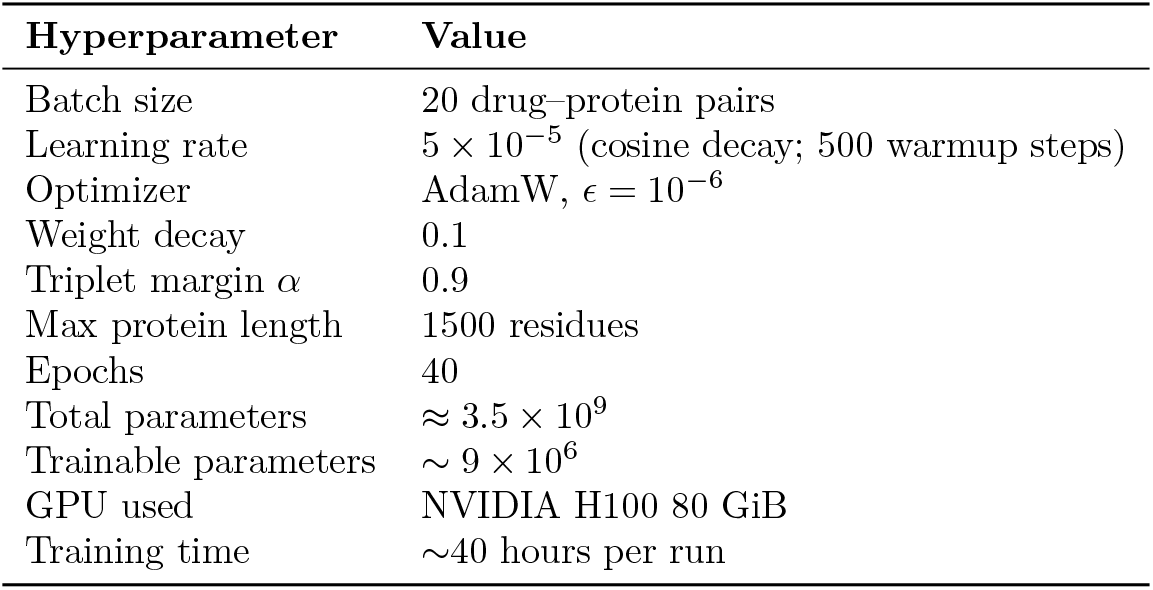
Training hyperparameters.

| Hyperparameter | Value |
| --- | --- |
| Batch size | 20 drug-protein pairs |
| Learning rate | $5 \times 10^{-5}$ (cosine decay; 500 warmup steps) |
| Optimizer | AdamW, $\epsilon = 10^{-6}$ |
| Weight decay | 0.1 |
| Triplet margin $\alpha$ | 0.9 |
| Max protein length | 1500 residues |
| Epochs | 40 |
| Total parameters | $\approx 3.5 \times 10^9$ |
| Trainable parameters | $\sim 9 \times 10^6$ |
| GPU used | NVIDIA H100 80 GiB |
| Training time | $\sim 40$ hours per run |

### A.3 Morgan fingerprint size ablation

To isolate the impact of ligand featurization, we train our model with Morgan fingerprints at multiple bit-lengths. Table 5 reports mean ± std PCC across five random seeds (best checkpoint per seed). Among the evaluated sizes, 2048-bit fingerprints provide the best average performance. We therefore adopt this setting as the default Morgan configuration for comparisons and downstream analyses, unless otherwise stated.

**Table 5:** Effect of Morgan fingerprint bit-length on BindingDB Patent (Pearson correlation, PCC). Results are mean ± std across 5 random seeds.

| Morgan FP size (bits) | PCC (mean ± std) |
| --- | --- |
| 1024 | $0.5710 \pm 0.0075$ |
| 2048 | $0.5701 \pm 0.0091$ |
| 4096 | $0.5627 \pm 0.0020$ |

### A.4 ADMET profile under different Morgan fingerprint sizes

To check whether the Morgan fingerprint resolution materially changes the physicochemical profile of retrieved ligands, we summarize major *predicted* ADMET properties for the final retrieved candidate sets under each fingerprint size. Table 6 reports mean ± std for molecular weight (MW), logP, HIA, hERG, QED, and Lipinski.

**Table 6:** Major predicted ADMET properties (mean ± std) for retrieved candidates under different Morgan fingerprint sizes.

| Morgan FP size (bits) | MW | logP | HIA | hERG | QED | Lipinski |
| --- | --- | --- | --- | --- | --- | --- |
| 1024 | $429.8 \pm 44.8$ | $3.46 \pm 1.20$ | $0.992 \pm 0.018$ | $0.776 \pm 0.149$ | $0.579 \pm 0.147$ | $3.69 \pm 0.45$ |
| 2048 | $465.9 \pm 38.5$ | $3.63 \pm 1.10$ | $0.997 \pm 0.005$ | $0.803 \pm 0.141$ | $0.598 \pm 0.115$ | $3.67 \pm 0.50$ |
| 4096 | $454.9 \pm 47.9$ | $3.09 \pm 1.05$ | $0.991 \pm 0.022$ | $0.799 \pm 0.144$ | $0.578 \pm 0.119$ | $3.65 \pm 0.44$ |

### A.5 Multi-objective focus: affinity with property preservation

In our multi-objective setting, the primary objective is *affinity*, i.e., we aim to increase predicted binding strength while remaining on the empirical molecule manifold. Nevermore therefore emphasizes affinity gains, but it does not do so by drifting into unrealistic edits. Two mechanisms keep the search property-aware: (i) a weighted ADMET penalty that discourages liability-increasing steps, and (ii) the projection step that maps each edited fingerprint back to a top-*K* neighborhood of *real* database molecules. Together, these constraints act as an invariance regularizer, yielding affinity improvements that remain consistent with desirable ADMET behavior. Figure 7 illustrates this behavior over 300 iterations: affinity increases while key endpoints remain stable or improve, with the EMA curve highlighting the underlying trend beneath projection-induced iteration-to-iteration jitter.

**Figure 7.**
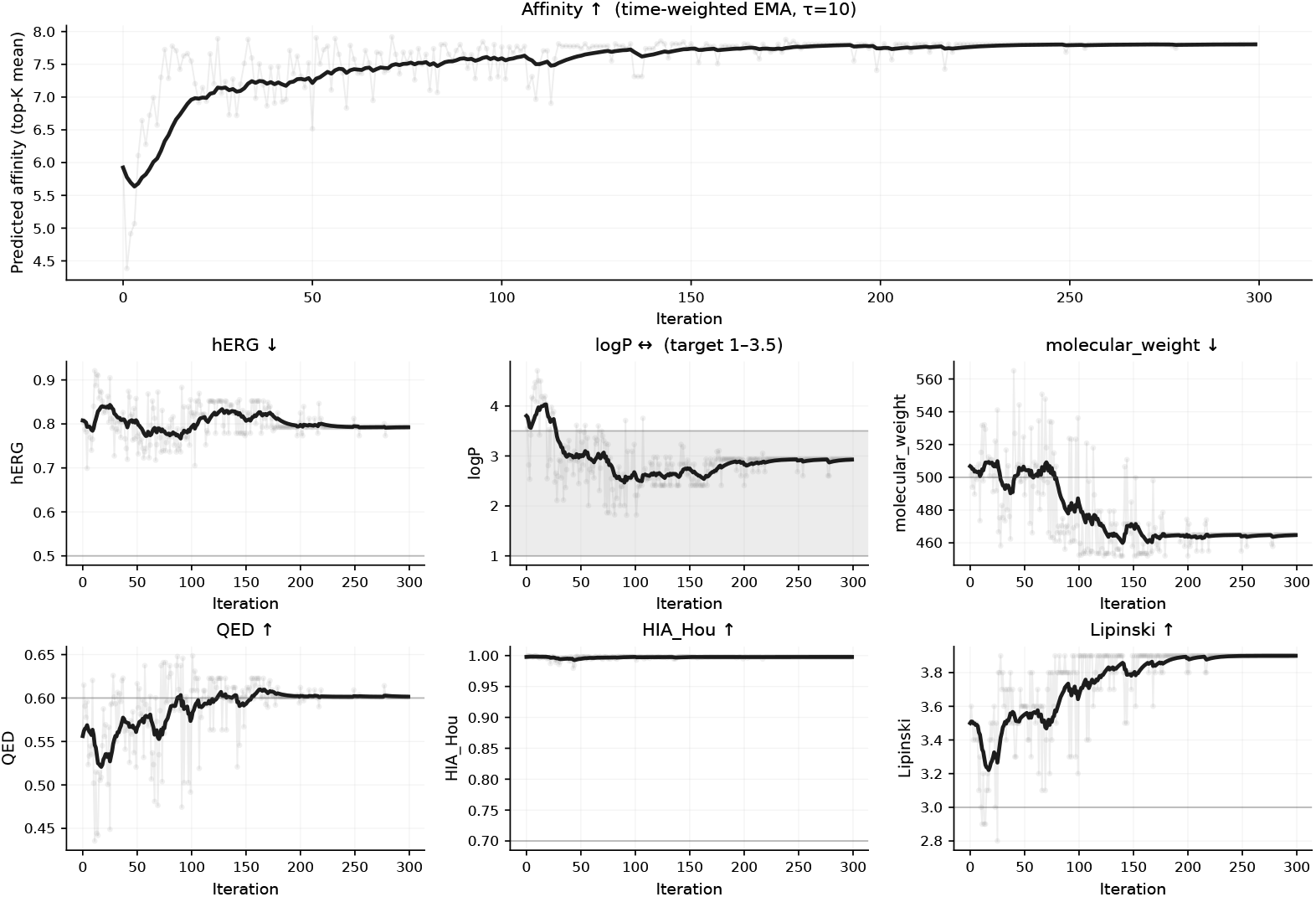
Nevermore optimization trajectories over 300 iterations. Top: predicted affinity statistic per iteration; bottom: corresponding ADMET endpoints. Light traces show raw per-iteration values and bold curves show a time-weighted EMA to emphasize trend.

#### Sensitivity to retrieval neighborhood size *k*

Figure 8 reports optimization trajectories for predicted affinity and developability proxies under different retrieval neighborhood sizes (*k* ∈ {1, 10, 100}). Small *k* (e.g., *k*=1) yields higher-variance trajectories because discrete fingerprint edits can change the identity of the single nearest neighbor abruptly, producing noisy objective feedback. Large *k* (e.g., *k*=100) smooths trajectories by averaging over a broader neighborhood, but can weaken the optimization signal by diluting the effect of sparse, integer edits. We therefore use *k*=10 as a practical trade-off that stabilizes optimization while preserving responsiveness to steering.

**Figure 8.**
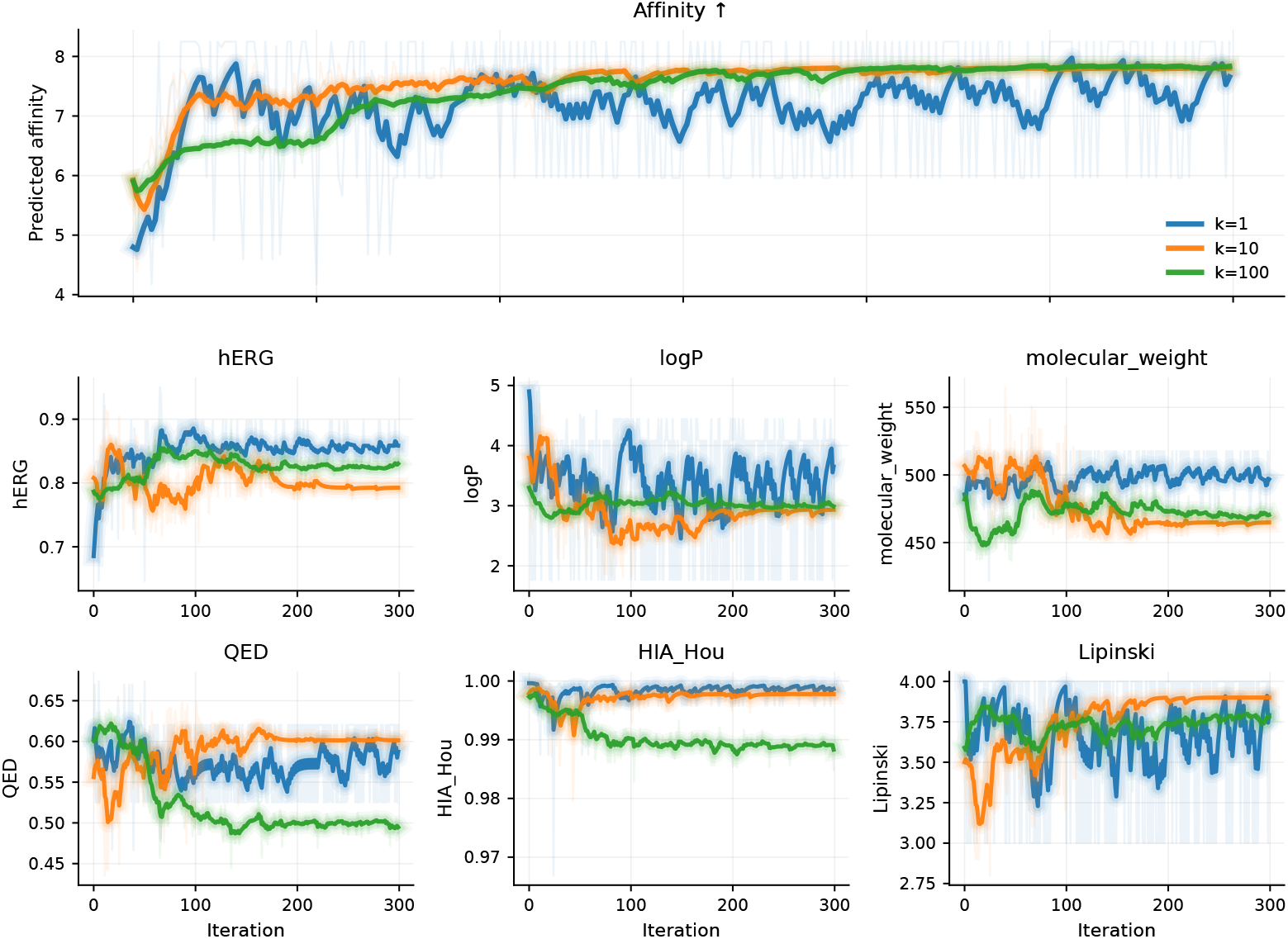
Sensitivity to retrieval neighborhood size *k*. Optimization trajectories for predicted affinity and developability proxies under different retrieval neighborhood sizes (*k* ∈ {1, 10, 100}). Small *k* (e.g., *k*=1) yields higher-variance trajectories because discrete fingerprint edits can change the identity of the nearest neighbor abruptly, producing noisy objective feedback. Large *k* (e.g., *k*=100) smooths trajectories by aggregating over a broader neighborhood, but can weaken the optimization signal by diluting the effect of sparse, integer edits. We therefore use *k*=10 as a practical trade-off that stabilizes optimization while preserving responsiveness to steering.

## Footnotes

1 https://huggingface.co/facebook/esm2_t12_35M_UR50D

